# Using Automated Detection and Classification of Interictal HFOs to Improve the Identification of Epileptogenic Zones in Preparation for Epilepsy Surgery

**DOI:** 10.1101/680280

**Authors:** Sina Farahmand, Tiwalade Sobayo, David J. Mogul

## Abstract

**Objective:** For more than 25 million drug-resistant epilepsy patients, surgical intervention aiming at resecting brain regions where seizures arise is often the only alternative therapy. However, the identification of this epileptogenic zone (EZ) is often imprecise which may affect post-surgical outcomes (PSOs). Interictal high-frequency oscillations (HFOs) have been revealed to be reliable biomarkers in delineating EZ. In this paper, an analytical methodology aiming at automated detection and classification of interictal HFOs is proposed to improve the identification of EZ. Furthermore, the detected high-rate HFO areas were compared with the seizure onset zones (SOZs) and resected areas to investigate their clinical relevance in predicting PSOs.

**Methods:** FIR band-pass filtering as well as a combination of time-series local energy, peak, and duration analysis were utilized to identify high-rate HFO areas in interictal, multi-channel intracranial electroencephalographic (iEEG) recordings. The detected HFOs were then classified into fast-ripple (FR), ripple (R), and fast-ripple concurrent with ripple (FRandR) events.

**Results:** The proposed method resulted in sensitivity of 91.08% and false discovery rate of 7.32%. Moreover, it was found that the detected HFO-FRandR areas in concordance with the SOZs would have better delineated the EZ for each patient, while limiting the area of the brain required to be resected.

**Conclusion:** Testing on a dataset of 20 patients has supported the feasibility of using this method to provide an automated algorithm to better delineate the EZ.

**Significance:** The proposed methodology may significantly improve the precision by which pathological brain tissue can be identified.

## I. Introduction

Epilepsy disease afflicts more than seventy million people worldwide [1]. In approximately one-third of the cases, antiepileptic medications fail to control seizures [2]. Epilepsy surgery is an alternative treatment for these drug-refractory patients, in which areas of the brain causing seizures are either resected or ablated [3].

Resecting the entire epileptogenic zone (EZ), the regions of the brain responsible for generating seizures, is the main goal of the surgical intervention for intractable seizures [4]. That is, precise resection of the entire EZ leads to successful post-surgical outcomes (PSOs) in which no future seizures should occur. The seizure onset zone (SOZ), the region of the brain where clinical seizures initiate, is thought to most closely localize the EZ. However, localization of the SOZ is often restricted due to the capability to record intracranial electro-encephalographic (iEEG) signals from a spatially limited brain area and the potential for rapid spread of seizure activity [5].

Recently, several studies have suggested that localized high frequency oscillations (HFOs) detected during interictal iEEG recordings are relevant biomarkers to localize the EZ [6]-[9]. It has also been reported that interictal HFOs are more abundant during non-rapid eye movement (non-REM) sleep compared to other stages of sleep [10]. HFOs are characterized as spontaneous, non-linear, non-stationary, and low-amplitude electrophysiological activity with a frequency bandwidth of 80-500 Hz [11]. In general, HFOs are subdivided with respect to their spectral content into ripple (80-250 Hz) and fast-ripple (250-500 Hz) bands [12].

HFO detection is conventionally performed using visual inspection of long hours of iEEG recordings. However, the aforementioned process is very tedious and time-consuming even for an expert epileptologist [13]. It also requires a great amount of concentration to eliminate either missing or false detection of any HFO events. Therefore, several automated HFO-detection methods have been proposed to overcome the aforementioned challenges.

Finite impulse response (FIR) or infinite impulse response (IIR) band-pass filtering of iEEG recordings, as well as using root-mean-squared (RMS) feature of the filtered signals, have been widely utilized to automatically detect HFO events [14]. The primary drawback of using this method is the high false discovery rate (FDR), which mainly stems from filtering of transient signals such as artifacts, spikes, and sharp waves without any superimposed HFOs [15]. That is, the FIR filtering of these broadband spectrum transient signals may generate oscillations in both ripple and fast-ripple bands, which represent spurious HFOs, that can be incorrectly attributed as true HFO events [15]. Therefore, it is of a crucial importance to combine FIR-filtering based methodologies with a FDR reduction process in order to separate and remove spurious HFOs from true HFO events.

Many time-frequency analyses such as Fourier, wavelet, and matching pursuit have been also used in detecting HFOs [14]-[17]. These methods, while convenient, assume a priori basis functions of the wideband electrophysiological signals such as cosine, sine, and different wavelet and Gaussian shapes. Therefore, they may fail to decompose signals into a set of narrowband components based on their intrinsic non-stationary and non-linear features.

Although several automated HFO detection methods have been proposed in the literature, the evidence for the correlation between the clinical relevance of the high-rate HFO areas in identifying the EZ and predicting the PSOs is rather poor [18]. That is, the application of using detected high-rate HFO areas as a resecting-map for epilepsy surgery is in an early stage of development and requires more investigation.

In this paper, an automated detection and classification of interictal HFOs is proposed to identify the high-rate HFO areas in multi-channel iEEG recordings. Moreover, different types of detected HFO areas were compared with the resected areas and SOZ to evaluate the relevance of areas with high-rate HFO activity in identifying the EZ and predicting PSOs.

The paper is organized as follows. Section II discusses the specifications of the adopted clinical dataset, along with the framework of the proposed HFO-detection method. In section III, performance analysis and experimental results of using the proposed HFO analysis to identify the EZ and predict PSOs are described. Section IV discusses the results, and Section IV provides conclusions to this research.

## II. Dataset and Methods

The conceptual framework of the proposed automated HFO detection and classification is illustrated in Fig. 1(a). Multi-channel iEEG data recorded from different brain regions of epilepsy patients were used as input to the analytical signal processing procedure. Following a pre-processing step on the iEEG data, the output signal was filtered within ripple and fast-ripple bandwidths in order to be used for HFO event detection. Next, a combination of signal processing procedures were performed in order to select HFO candidates from these oscillators. Following an automated FDR reduction process, the final HFO events were obtained and saved in the HFO database. Fig. 1(b) exhibits the framework for evaluating clinical relevance of the detected final HFOs, in identifying the EZ and predicting PSOs, using both rate thresholding and resection ratio (RR) analysis.

**Fig. 1.**
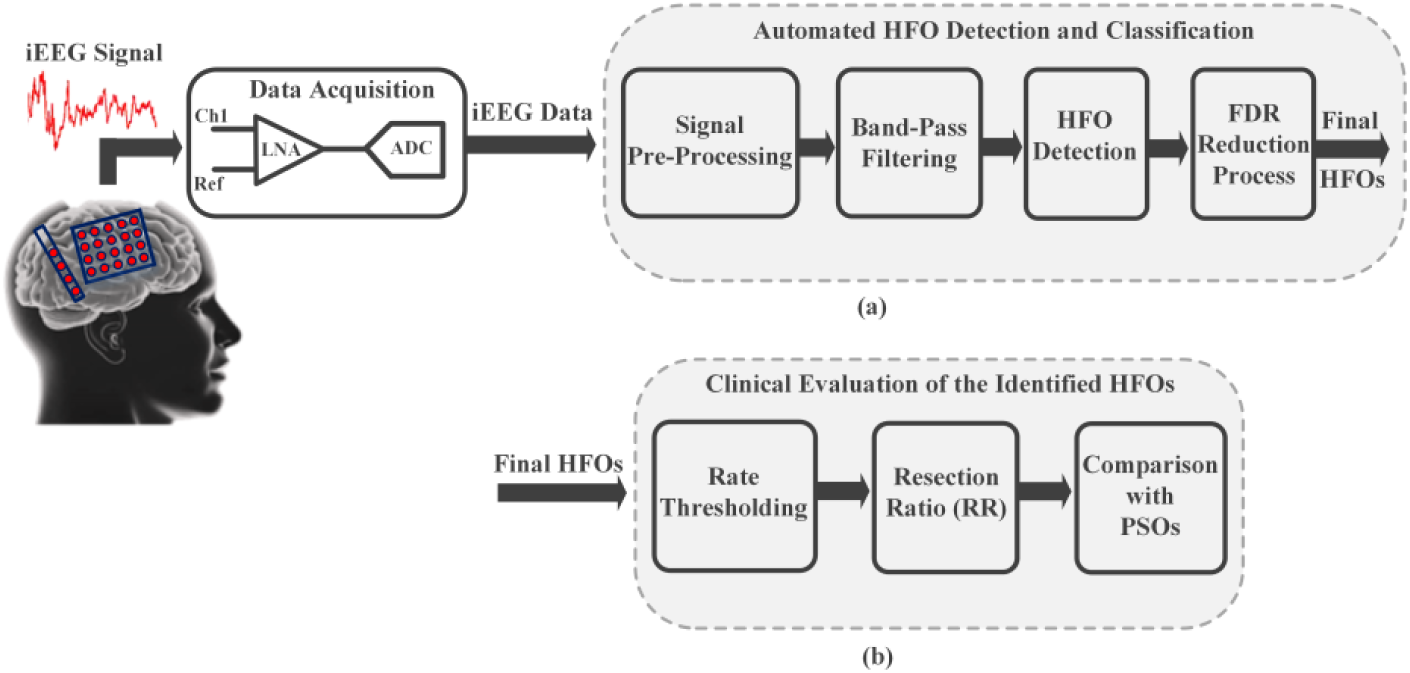
Framework of the proposed HFO detection and classification methodology in identifying the EZ and predicting PSOs. (a) Automated HFO detection and classification. (b) Clinical evaluation of the identified HFO areas.

### A. Patients and iEEG Dataset

In this study, invasive, multi-channel iEEG data recorded from 20 patients who afterward underwent epilepsy surgery, at the Neurosurgery Department of the University Hospital of Zurich, Switzerland were used [19]. The iEEG recordings were performed using subdural strip, grid, as well as depth electrodes. For each epilepsy patient, up to six intervals, each containing five minutes of interictal slow-wave sleep, were selected for the analysis. In this stage of sleep, interictal HFO activities are abundant compared to other sleep stages [10]. Long-term data acquisition was carried out using a Neuralynx system with a sampling frequency of 4000 Hz, which was later down-sampled to 2000 Hz for HFO analysis, and a 0.5-1000 Hz band-pass filtering. Furthermore, a digital 60 Hz notch filter was used to eliminate line noise. Additional clinical information regarding the 20 analyzed patients and dataset are provided in Table I. For each epilepsy patient, the channels within the SOZ were assigned by epileptologists through identification of the region most proximal to iEEG recording sites where ictal discharges were first observed.

**TABLE I.** Patient Clinical Information

| Patient ID | Age(y) / Sex | Epilepsy | Outcome (ILAE) | Types of electrodes |
| --- | --- | --- | --- | --- |
| Patient-1 | 25 / M | TLE | 1 | 1 strip 4x1, 1 strip 6x1, 5 depth |
| Patient-2 | 33 / M | TLE | 1 | 8 depth |
| Patient-3 | 20 / F | TLE | 1 | 5 depth |
| Patient-4 | 20 / F | TLE | 1 | 8 depth |
| Patient-5 | 40 / M | TLE | 1 | 8 depth |
| Patient-6 | 48 / M | TLE | 1 | 8 depth |
| Patient-7 | 25 / M | TLE | 3 | 8 depth |
| Patient-8 | 21 / F | TLE | 3 | 8 depth |
| Patient-9 | 52 / M | TLE | 5 | 8 depth |
| Patient-10 | 37 / M | ETE | 1 | 1 grid 8x4, 2 strips 4x1 |
| Patient-11 | 36 / M | ETE | 1 | 1 grid 8x8, 1 depth |
| Patient-12 | 49 / M | ETE | 1 | 1 grid 8x4, 1 depth |
| Patient-13 | 17 / M | ETE | 1 | 1 grid 8x8, 1 depth |
| Patient-14 | 46 / F | ETE | 1 | 2 grids 8x2, 1 strip 6x1, 1 strip 4x1, 1 depth |
| Patient-15 | 31 / F | ETE | 1 | 1 grid 8x4, 2 strips 4x1 |
| Patient-16 | 17 / F | ETE | 1 | 1 grid 8x4, 1 depth |
| Patient-17 | 30 / M | ETE | 5 | 2 grids 8x2, 1 depth |
| Patient-18 | 40 / M | ETE | 5 | 2 strips 6x1, 1 depth |
| Patient-19 | 38 / M | ETE | 6 | 1 grid 8x4, 1 grid 8x2 |
| Patient-20 | 17 / M | ETE | 5 | 1 grid 8x2 |

M= male; F= female; ILAE= international league against epilepsy, TLE= temporal lobe epilepsy; ETE= extratemporal lobe epilepsy; grid= grid electrodes; depth= depth electrodes.

### B. Signal Pre-Processing

The multi-channel iEEG signals were recorded based on a common intracranial reference. As the first stage of the signal pre-processing, bipolar derivations from iEEG signals were measured by taking the voltage difference between adjacent channels. This is done to eliminate the confounding effects of both common reference signal and volume conduction in the HFO events detection [20], [21]. Next, the bipolar-derivate signals were divided in consecutive, non-overlapping, 1-s windows. The second stage of the signal pre-processing was to eliminate the edge effect problem related to the data segmentation prior to executing the FIR band-pass filtering on each 1-s window. In order to do that, 500 samples from neighboring segments were incorporated to both ends of each 1-s data segment. However, after performing the FIR filtering, only the output corresponding to the original data segment were retained.

### C. FIR Band-Pass Filtering

FIR band-pass filtering was carried out on each bipolar-derivate iEEG data-segment in the ripple and fast-ripple bands in order to prepare data for HFO detection. In the fast-ripple band, each data-segment was filtered using a FIR equiripple filter with a pass-band between 250-490 Hz and the stop-band of 240 Hz and 500 Hz with a 60 dB stop-band attenuation. In the ripple band, the same FIR filter-type was used with a pass-band between 80-240 Hz and the stop-band of 70 Hz and 250 Hz with a 60 dB stop-band attenuation. It should be noted that a FIR filter was preferred over IIR filter due to its linear phase properties as well as its less ringing tendency especially when dealing with a high attenuation level [15]. Each data-segment was also band-pass filtered in the gamma band merely for the FDR-reduction purpose as described later in this paper. In the gamma band, the FIR equiripple filter was used with a pass-band between 40-70 Hz and the stop-band of 30 Hz and 80 Hz with a 60 dB stop-band attenuation. Fig. 2(a) illustrates a 1-s bipolar-derivate iEEG data-segment containing ripple activity along with its FIR filtered version in the ripple band. Fig. 2(b) also exhibits a 1-s iEEG data-segment containing fast-ripple activity along with its FIR band-pass filtered version in the fast-ripple band.

**Fig. 2.**
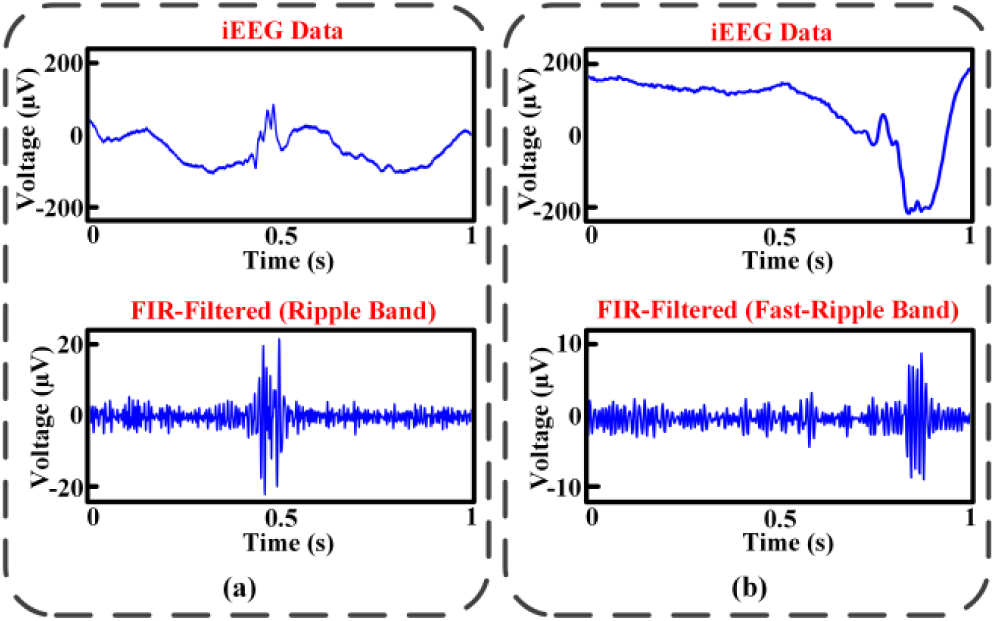
FIR band-pass filtering of a 1-s, bipolar-derivate iEEG signal. (a) 1-s bipolar-derivate iEEG data-segment containing ripple activity along with its FIR filtered version in the ripple band. (b) 1-s iEEG data-segment containing fast-ripple activity along with its FIR band-pass filtered version in the fast-ripple band.

### D. HFO Detection

Following the FIR band-pass filtering on the bipolar-derivate iEEG data and extracting filter outputs or neuronal oscillators in the ripple and fast-ripple bands, their root-mean-squared (RMS) amplitude were measured individually using a 2-ms sliding window in order to determine the local energy of the oscillators. Next, an RMS-threshold value was determined using the whole 5-min of each interval for both the ripple and fast-ripple filtered signals respectively. It should be noted that the RMS-thresholds were later optimized to maximize the sensitivity of the automated HFO detector while minimizing its FDR, which are described in detail in the results section. For each filtered signal, successive RMS values that exceed its corresponding RMS-threshold longer than 6-ms in duration were considered as putative HFOs and their onset and offset were marked and saved in the database. Moreover, the consecutive putative HFO events detected in the fast-ripple band, HFO-FR, as well as the ones detected in the ripple-band, HFO-R, separated by less than 10-ms were combined as one event respectively. The detected events within each band were then rectified and those with at least six peaks above a peak-threshold were classified as HFO-FR and HFO-R candidates. The peak-threshold was set to (µ + 3σ) of the filter output’s instantaneous amplitude using the whole 5-min of each interval. In addition, the HFO-FR candidates that occur concurrently with HFO-R candidates were classified as HFO-FRandR candidates.

### E. FDR Reduction Process

FIR band-pass filtering of high-energy spikes, artifacts, and sharp waves without superimposed HFOs is sensitive to the misclassification of the resultant HFOs as true HFOs.

In this study, the time-frequency properties of spikes along with a concurrent multi frequency-band analysis were utilized to significantly reduce the sensitivity of the proposed method to the misclassification of high-energy spikes as true HFOs. This process is described for monophasic and biphasic spikes in Fig. 3 and Fig. 4 respectively. Fig. 3(a) illustrates an intracranially-recorded monophasic spike, which comprises a relatively wide base and a pointed peak. Based on the time-frequency analysis using Fourier transform, the pointed peak of spike is made of high-frequency components while its base is constructed using lower-frequency components. Moreover, the temporal extent of high-frequency components is smaller than that of the lower-frequency ones. It should be noted that for the purpose of FDR-reduction, the same procedure for detecting the HFO-FR and HFO-R candidates, as mentioned in the last section, were applied to the filtered output of bipolar-derivate iEEG data-segments in the gamma band. Fig. 3(b) exhibits the signals resulted from filtering of the spike in the fast-ripple, ripple, and gamma bands. Fig. 3(c) shows the local energy of each FIR-filtered signal, measured using RMS values of their instantaneous amplitudes, over a 2-ms sliding window. The purple dashed lines denote the RMS-thresholds measured for each FIR-filtered signal over the entire 5-min of the interval in which the spike occurred. Furthermore, the green dashed lines indicate the onset and offset time when the local energy, for each filtered signal, exceeds its corresponding RMS-thresholds. The detected events in both ripple and gamma bands must individually satisfy their peak, energy, and duration criteria, mentioned in the last section, in order to be enrolled in spike rejection. As it is demonstrated in Fig. 3(c), the local energy of the gamma band encompasses the one in the ripple band considering both timing and energy intensity. It should be noted that the HFO-FR event, detected in its corresponding FIR-filtered signal, failed to exceed the RMS-threshold for more than 6-ms. Fig. 4 shows a synthesized biphasic spike along with its FIR-filtered signals in gamma, ripple, and fast-ripple bands and local energy characteristics. The local energy of the gamma band embraces the ones in the ripple and fast-ripple bands, and the local energy in the ripple band encompasses the one in the fast-ripple band. Moreover, the detected HFO-FR and HFO-R events satisfied all the energy, peak, and duration criteria.

**Fig. 3.**
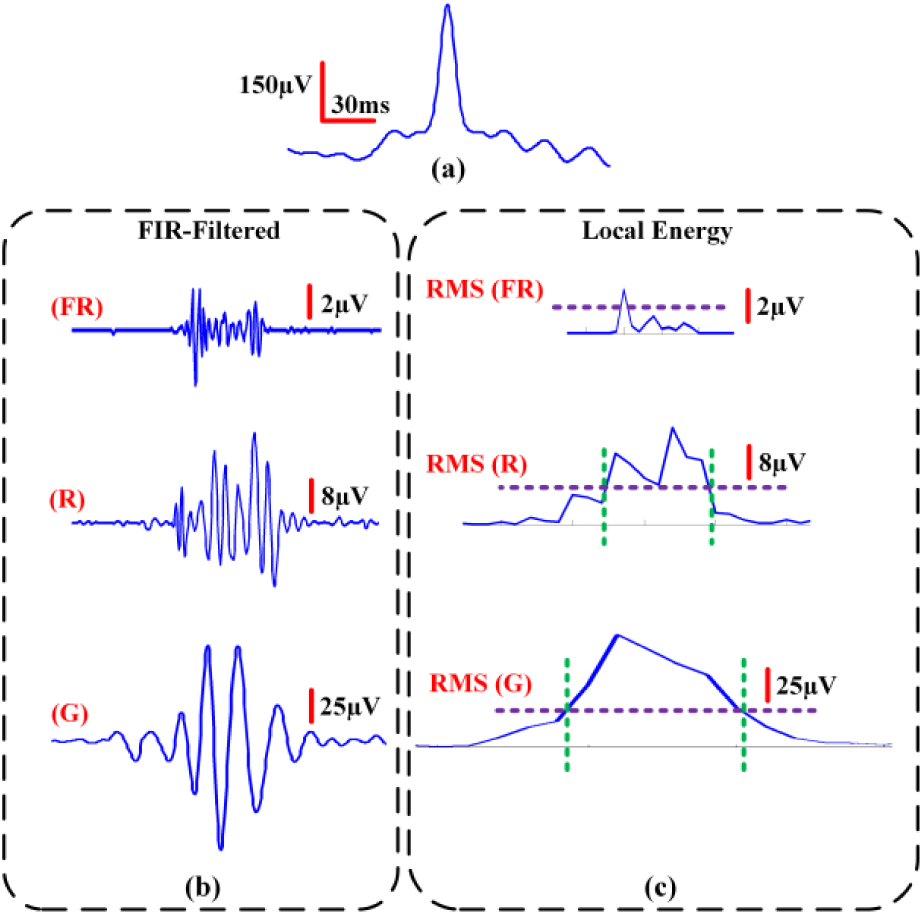
FDR reduction process for a monophasic spike. (a) An intra-cranially recorded monophasic spike used for FDR reduction process. (b) Output signals resulted from FIR band-pass filtering of the monophasic spike in the fast-ripple (FR), ripple (R), and gamma (G) bands. (c) Local energy of each FIR-filtered signal, measured using RMS values of their instantaneous amplitudes, over a 2-ms sliding window. The purple dashed lines indicate the RMS-threshold that were independently measured for each FIR-filtered signal over the entire 5-min of the interval in which spike occurred. The green dashed lines denote the onset and offset time when the local energy, for each FIR-filtered signal, exceeds its corresponding RMS-threshold. The green dashed lines were not shown for the local energy in the fast-ripple band because it failed to exceed its RMS-threshold for more than 6-ms.

**Fig. 4.**
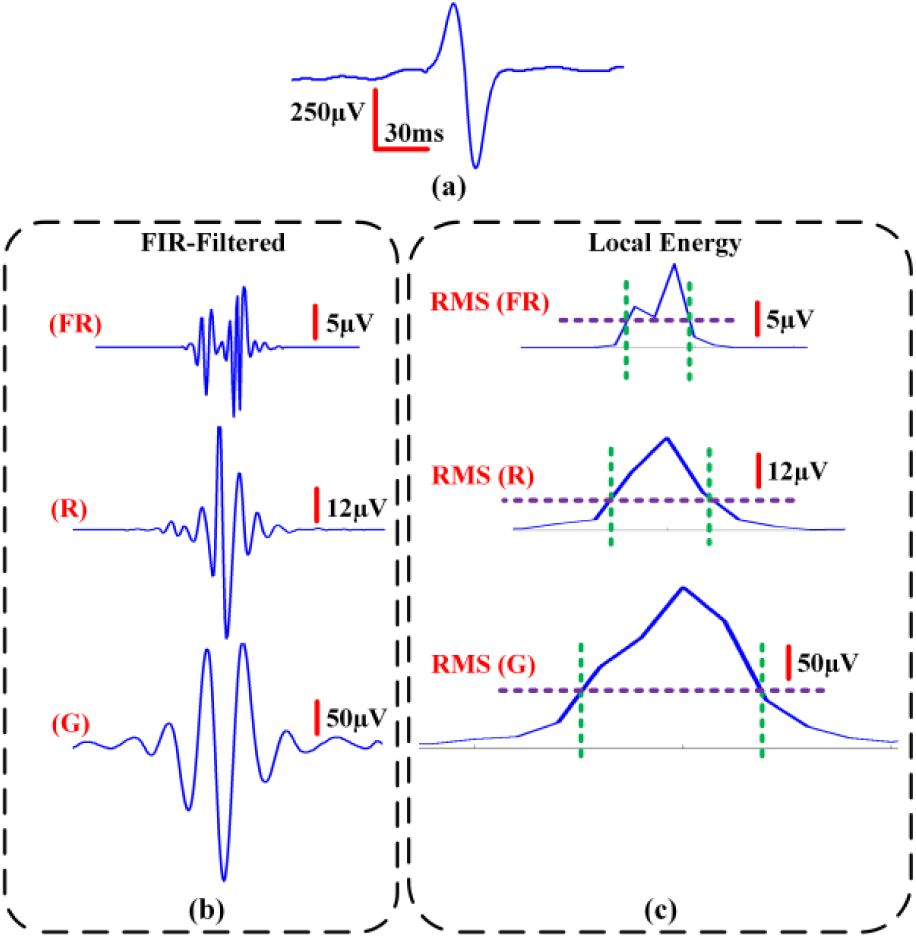
FDR reduction process for a biphasic spike. (a) A simulated biphasic spike used for FDR reduction process. (b) Output signals resulted from FIR band-pass filtering of the biphasic spike in the fast-ripple (FR), ripple (R), and gamma (G) bands. (C) Local energy of each FIR-filtered signal, measured using RMS values of their instantaneous amplitudes, over a 2-ms sliding window. The purple dashed lines indicate the RMS-threshold that were independently measured for each FIR-filtered signal over the entire 5-min of the interval, in which spike occurred. The green dashed lines denote the onset and offset time when the local energy, for each FIR-filtered signal, exceeds its corresponding RMS-threshold.

The extracted time-frequency features of spikes without HFOs were tested and validated using the visually detected spikes within 2-h of the iEEG dataset. These features were then used in the proposed classifier in order to discriminate spurious HFOs from true HFOs. These features are outlined as follows:

1. RMS(G) onset time < RMS(R) onset time < RMS(FR) onset time*
2. RMS(FR) offset time* < RMS(R) offset time < RMS(G) offset time
3. RMS(FR) energy* < RMS(R) energy < RMS(G) energy

* The FR-related criteria can be ignored in case the extracted FIR-filtered signal(FR) fails to satisfy the HFO-FR candidate conditions.

The RMS(x) energy indicates the measured RMS values of the FIR-filtered signal(x), where *x* ∈{‘*FR*’,‘*R*’,‘*G*’}, during the time that it exceeds its corresponding RMS-threshold.

The detected HFO-candidates, from the previous stage, that fulfil all the aforementioned three criteria were removed from the HFO database and the remaining ones were considered and classified as final HFOs; namely, final HFO-FR, final HFO-R, and final HFO-FRandR.

### F. Visual Inspection and Performance Analysis

Visual inspection of 2-h of bipolar-derivate, interictal iEEG signals were carried out independently by two reviewers trained in electrophysiology and HFO analysis in order to analyze the performance of the proposed HFO detector. The zoomed-in interictal iEEG signals, 1-s/plot, were provided in the first trace while the FIR band-pass filtered signals in the ripple and fast-ripple bands were provided simultaneously in the second and third trace. Besides, the FIR filtered signals in the fast-ripple and ripple bands were viewed at a higher gain due to their lower amplitudes compared to the unfiltered input signal. Reviewers used the FIR-filtered data in order to verify their visually detected HFOs. Next, they classified and labeled the visually marked HFO events from the iEEG data as HFO-FR and HFO-R. They also marked spikes without HFOs on the iEEG data merely to verify the FDR-reduction process as described in the last section. All the visually detected HFO events that were jointly marked by both reviewers, in ripple and fast-ripple bands, were considered as a gold standard to quantitatively evaluate the performance of the automated HFO detector using sensitivity and FDR parameters. The sensitivity parameter measures the proportion of HFO events, detected using our methodology, overlapping with the visually detected ones and is defined as follows:

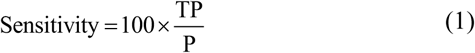

in which *TP* indicates the number of correctly detected HFO events and *P* is the total number of visually detected HFOs. The FDR parameter calculates the incorrectly detected HFO events and is described by equation (4).

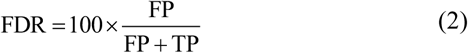

where *FP* represents the number of incorrectly detected HFO events such as interictal spikes, artifacts, and sharp waves.

### G. Clinical Evaluation of the Identified HFOs

In order to assess the clinical relevance of the detected final HFOs in different classes, in detecting the EZ and predicting PSOs, the rate of their HFO activities were computed for each 5-minutes interval. Next, the median rate of the intervals was chosen to represent the HFO rate of a particular channel for each patient. Finally, a rate threshold was measured separately for different classes of final HFOs using Kittler’s method to identify channels with high HFO rates [22]-[25]. Kittler’s algorithm is a histogram-based thresholding method that approximates the histogram as a bimodal distribution and finds the cutoff point to separate it. Due to the small sample size, the bootstrap method was used to reliably estimate the rate threshold [26]. One thousand bootstrap samples were generated, with replacement from the original dataset, containing the median rates of each channel. Kittler’s threshold was measured for each of the 1000 bootstrap samples and the mean value of the obtained 1000 thresholds was chosen as the final threshold to separate the high-rate HFO channels, defined as HFO areas, from the ones with low rates of HFOs. In addition to Kittler’s method, we also tried to separate channels with high HFO rates using Tukey’s upper fence and the 95^th^ percentile. Kittler’s method provided results with the highest sensitivity with the resected area.

In order to assess the clinical relevance of the detected HFO areas, in all three classes, in predicting PSOs the resection ratio (RR) of the HFO areas is described as follows:

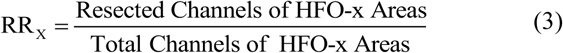

in which subscript *x* indicates different HFO classes; namely, FR, R, and FRandR. The RR has values from zero to one.

## III. Experimental Results

In this section, the performance of the proposed automated HFO detector is assessed using the jointly marked HFO events by both reviewers, before and after the FDR-reduction process. Moreover, the identified HFO areas were compared with the SOZ and resected areas for each patient to investigate their clinical relevance in detecting the EZ and predicting PSOs.

### A. Performance Results

In the training stage of the HFO-events detection, the RMS-threshold was optimized individually for the FIR-filtered signals in the ripple and fast-ripple bands to enhance the detection sensitivity while reducing the FDR. During this stage, the peak-threshold was kept fixed and equal to the (µ + 3σ) of the filter output’s instantaneous amplitude using the whole five minutes of each interval. Next, the RMS-threshold was varied according to the formula (µ + kσ), where *µ* is the mean of the RMS values over the 5-min interval, *σ* is their standard deviation, and *k* ranged from 0 to 10 with 0.5 increments. A maximum value of 10 was selected for k to guarantee that at least 99% of the RMS values lie below the RMS-threshold according to Chebyshev’s inequality [27]. To find the optimum RMS-threshold for HFO detection, the ROC curve was used to plot the sensitivity as a function of FDR, when varying RMS-threshold [28]. Fig. 5 illustrates the ROC curves for HFO-FR, blue curve, and HFO-R, orange curve, events. The optimum RMS-threshold value for both HFO-FR and HFO-R, where the discrimination between the sensitivity and FDR is maximum, were found equal to (µ + 5σ). The obtained optimum RMS-threshold was consistent among all intervals within the 2-h test data. The automated HFO detector was tuned to that obtained, optimum RMS-threshold for HFO detection, classification, and FDR-reduction process. The sensitivity and FDR of the automated HFO detector prior and after the FDR-reduction process are provided in Table II. The reported sensitivity and FDR values of the HFO detector were measured with respect to the jointly marked HFO events by reviewers.

**TABLE II.**
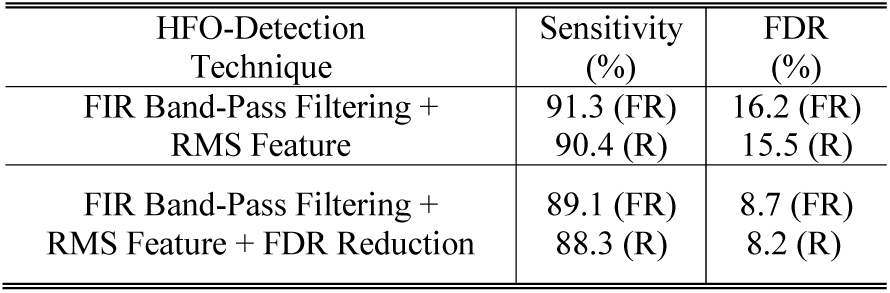
Performance of the Proposed HFO-Detection Method

| HFO-Detection Technique | Sensitivity (%) | FDR (%) |
| --- | --- | --- |
| FIR Band-Pass Filtering + RMS Feature | 91.3 (FR)<br>90.4 (R) | 16.2 (FR)<br>15.5 (R) |
| FIR Band-Pass Filtering + RMS Feature + FDR Reduction | 89.1 (FR)<br>88.3 (R) | 8.7 (FR)<br>8.2 (R) |

**Fig. 5.**
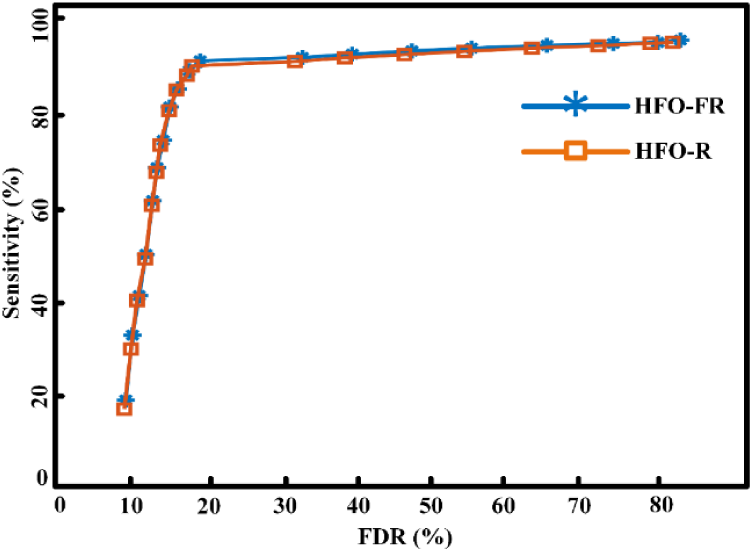
The simulated ROC curves for both HFO-FR, blue curve, and HFO-R, orange curve, events. It should be noted that the RMS-threshold was optimized individually for the FIR-filtered signals in the ripple and fast-ripple bands.

### B. HFO-Area Identification

In this section, the identification of channels with high-rate HFO activities, HFO-areas, from 24 bipolar-derivate channels is described. For simplicity, only the detected HFO-FR areas from all six intervals is demonstrated. Fig. 6(a) illustrates the histogram of the detected HFO-FR events across all channels. The x-axis indicates the median of the HFO-FR rate values from all six intervals, each containing 5 min. of iEEG data. A rate-threshold, denoted by a purple dashed line, equal to 8, was measured based on Kittler’s method for all channels. Fig. 6(b) exhibits the channels in which the median of their HFO-FR rate values, from all six intervals, exceeded the measured rate-threshold.

**Fig. 6.**
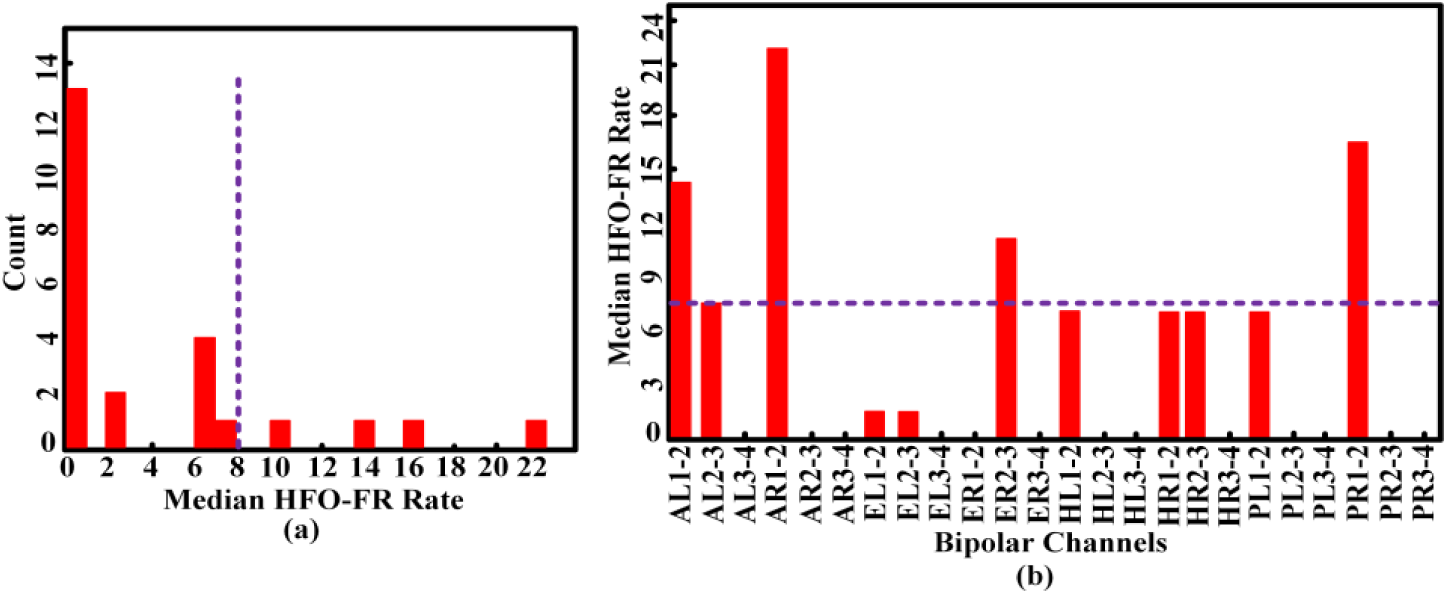
Identification of channels with high-rate HFO-FR activities. (a) Histogram of the detected HFO-FR events across all 24 bipolar-derivate channels. The x-axis denotes the median of the HFO-FR rate values from all six intervals. (b) The identified channels with median HFO-FR rate, measured from all six intervals, above the rate threshold.

### C. Clinical Assessment of the Identified HFO-Areas

For all the patients reported in Table I, seizure outcome of the resecting surgery was evaluated in subsequent visits and then classified according to the International League Against Epilepsy (ILAE) scale [29]. For instance, Patient-1, Patient-2, and Patient-3 were reported seizure-free after surgery (ILAE Class 1, Table I). However, Patient-9 (ILAE Class 3, Table I) Patient-17, and Patient-18 (ILAE Class 5, Table I) still had recurrent seizures. For each patient, a summary of the resulted RR values for different HFO classes; namely, HFO-FR, HFO-R, and HFO-FRandR is reported in Table III. It was found that the identified HFO-FRandR areas better predicted PSOs relative to the other two classes. The *RR*_*(FRandR)*_ values equal to 1 indicated that all the identified HFO-FRandR areas, using the proposed method, were within the resected areas of brain. Consistent with the notion that HFO-generating areas are a suitable biomarker for the EZ, the identified HFO-FRandR channels in the patients overlapped well with the SOZ. In 17 out of 20 patients, the HFO-FRandR area formed a subset of the SOZ, while the remaining three patients had an overlap greater than 50%. It should be noted that the identified HFO areas for FR, R, and FRandR can be distinct. In particular, as shown for Patient-8 and Patient-9, it is possible for the HFO-R and HFO-FR areas to be fully resected, but not the identified high-rate HFO-FRandR channels or HFO-FRandR areas. That is, PSOs of epilepsy patients showed a high specificity to the identified HFO-FRandR areas.

**TABLE III.** Resection Ratio (RR) for the Identified HFO Areas in Different HFO Classes

| Patient ID | Outcome (ILAE) | RR<sub>(FR)</sub> | RR<sub>(R)</sub> | RR<sub>(FRandR)</sub> |
| --- | --- | --- | --- | --- |
| Patient-1 | 1 | 1.0 | 1.0 | 1.0 |
| Patient-2 | 1 | 0.67 | 0.67 | 1.0 |
| Patient-3 | 1 | 0.5 | 0.4 | 1.0 |
| Patient-4 | 1 | 1.0 | 1.0 | 1.0 |
| Patient-5 | 1 | 0.67 | 0.75 | 1.0 |
| Patient-6 | 1 | 1.0 | 1.0 | 1.0 |
| Patient-7 | 3 | 0.33 | 0.25 | 0.0 |
| Patient-8 | 3 | 1.0 | 1.0 | 0.33 |
| Patient-9 | 5 | 1.0 | 1.0 | 0.5 |
| Patient-10 | 1 | 0.75 | 0.63 | 1.0 |
| Patient-11 | 1 | 0.88 | 0.75 | 1.0 |
| Patient-12 | 1 | 0.75 | 0.5 | 1.0 |
| Patient-13 | 1 | 1.0 | 0.5 | 1.0 |
| Patient-14 | 1 | 0.5 | 0.33 | 1.0 |
| Patient-15 | 1 | 0.8 | 0.6 | 1.0 |
| Patient-16 | 1 | 0.5 | 0.14 | 1.0 |
| Patient-17 | 5 | 0.63 | 0.5 | 0.6 |
| Patient-18 | 5 | 0.33 | 0.5 | 0.0 |
| Patient-19 | 6 | 0.88 | 0.76 | 0.5 |
| Patient-20 | 5 | 0.0 | 0.0 | 0.0 |

## IV. Discussion

The primary aim of this study was to develop an automated HFO detection and classification methodology to improve the identification of the EZ prior to epilepsy surgery. The clinical relevance of different classes of identified HFO areas; namely, HFO-FR, HFO-R, and HFO-FRandR in predicting PSOs was investigated using rate thresholding and RR. A combination of FIR band-pass filtering and FDR reduction process was utilized to not only reliably detect HFO events, but also to decrease the sensitivity of the proposed method to the misclassification of high-energy transients such as sharp waves and spikes as true HFOs.

The FDR reduction process was not only capable of removing the spurious HFOs created from the FIR-filtering of spikes without HFOs, it was also sensitive to detecting HFO events superimposed on spikes or sharp waves. This mainly stemmed from the following two reasons. First, HFO events can take place at any portion of spikes; therefore, the first and second spike-rejection criteria corresponding to onset and/or offset times are not necessarily fulfilled. Second, HFO-FR and/or HFO-R superimposed on spikes can enhance their corresponding local energy level that may contradict the third spike-rejection criteria.

Patient-16 (ILAE Class 1, Table I) and Patient-17 (ILAE Class 5, Table I) were selected as case studies to discuss and visually demonstrate the process for investigating the clinical relevance of the identified HFO-FRandR areas in identifying the EZ and predicting PSOs. Resecting the HFO-FRandR areas was found to be better than the HFO-FR areas and the HFO-R areas at predicting PSOs on the group level. This finding is consistent with other studies [30], [31]. Fig. 7(a)-(b) show the anatomical position of the implanted electrodes in Patients-16 and Patient-17 as well as their SOZ, determined by clinicians, as electrode sites with blue ×. Moreover, the HFO-FRandR areas obtained using our automated methodology are presented with electrode contacts filled with red. The identified HFO-FRandR areas, in both patients, overlapped well with their SOZ. Furthermore, they are smaller than the annotated SOZ and form a subset of it. Our analysis suggests the EZ in these patients might have been significantly smaller than the SOZ identified by clinicians. Therefore, the proposed HFO analysis in these epilepsy patients might have significantly improved the surgical planning that defined the resected area. Moreover, Patient-16 who was seizure-free after epilepsy surgery had his identified HFO-FRandR area fully resected, but not the entire annotated SOZ. On the other hand, Patient-17 did not have a good PSO despite the resection of most of the SOZ. For Patient-17, a large portion of the right frontal cortex was resected. Although parts of the HFO-FRandR areas identified using the proposed HFO detection and classification methodology were included in the resection, its resection was not complete.

**Fig. 7.**
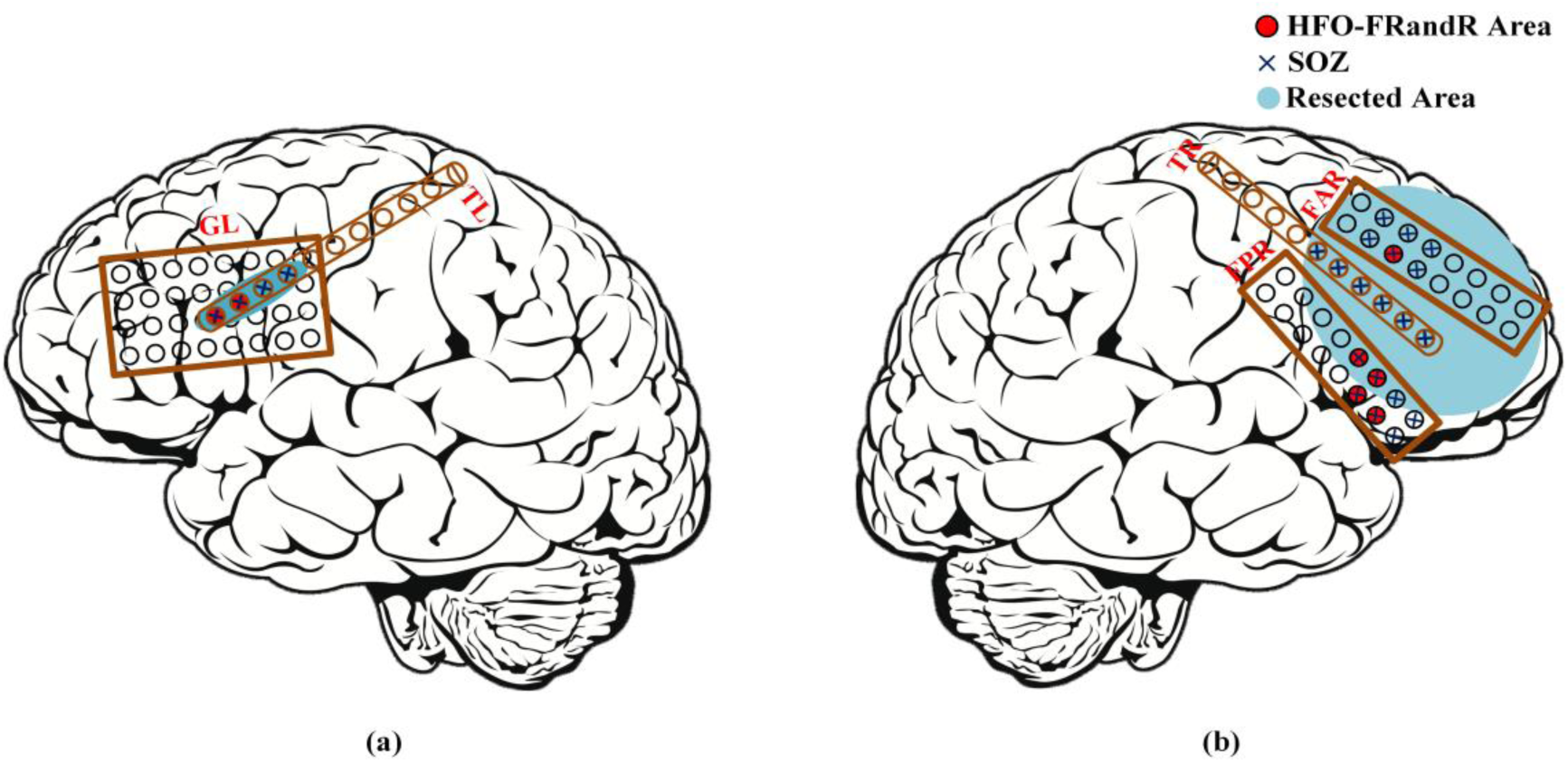
The HFO-FRandR areas identified in Patient-16 and Patient-17 with extratemporal lobe epilepsy. (a) Anatomical position of the implanted grid and depth electrodes for Patient-16. (b) Anatomical position of the implanted grid and depth electrodes for Patient-17. The SOZ identified by clinician are denoted as electrode contacts with the blue ×. The HFO-FRandR areas detected by the proposed analysis is denoted by contacts filled with red color. The resected areas during epilepsy surgery are represented by blue ellipse.

Our results suggest that the detected HFO-FRandR areas in concordance with the SOZ would have better delineated the EZ, while limiting the area of the brain required to be resected. This might have improved the PSO for Patient-17 with less destruction of brain tissue and therefore, fewer side effects.

Although this protocol will need to be assessed further using additional epilepsy patients, it provides an analytical protocol that may not only increase surgical efficacy for patients, but it may also minimize neurological deficits that may arise as a result of this surgery.

## V. Conclusion

In this study, an automated HFO detection and classification methodology is proposed to improve the localization of the EZ in patients with drug-resistant epilepsy using interictal iEEG recordings. Different types of HFOs; namely, HFO-FR, HFO-R, and HFO-FRandR were detected and classified using FIR band-pass filtering in the ripple and fast-ripple bands along with a HFO detection process. Moreover, the spurious HFOs caused by FIR-filtering of spikes without HFOs were removed from the final HFO database using the FDR reduction process. Next, the high-rate HFO channels were identified in all three classes to evaluate the clinical relevance of them in localizing the EZ and predicting PSOs. Our results suggested that patients who had their HFO-FRandR areas fully resected ended up seizure-free while patients with HFO-FRandR areas that were not fully resected still had recurrent seizures.

Testing on a preliminary dataset of 20 epilepsy patients has supported the feasibility of using our method to provide an automated algorithm that can be used in concordance with the SOZ to better delineate the EZ. This could potentially lead to an improvement in the surgical planning of the resected area.

## Acknowledgment

The authors would like to thank the lab of Dr. Johannes Sarnthein at the University of Zurich, and colleagues at the Swiss Epilepsy Center, for providing us the clinical dataset and information regarding SOZs and resected areas.

